# FMRP Links Optimal Codons to mRNA stability in Neurons

**DOI:** 10.1101/801449

**Authors:** Huan Shu, Elisa Donnard, Botao Liu, Ruijia Wang, Joel D. Richter

## Abstract

Fragile X syndrome (FXS) is caused by inactivation of the *FMR1* gene and loss of encoded FMRP, an RNA binding protein that represses translation of some of its target transcripts. Here we use ribosome profiling and RNA-seq to investigate the dysregulation of translation in the mouse brain cortex. We find that most changes in ribosome occupancy on hundreds of mRNAs are largely driven by dysregulation in transcript abundance. Many downregulated mRNAs, which are mostly responsible for neuronal and synaptic functions, are highly enriched for FMRP binding targets. RNA metabolic labeling demonstrates that in FMRP-deficient cortical neurons, mRNA downregulation is caused by elevated degradation, and is correlated with codon optimality. Moreover, FMRP preferentially binds mRNAs with optimal codons, suggesting that it stabilizes such transcripts through direct interactions via the translational machinery. Finally, we show that the paradigm of genetic rescue of FXS-like phenotypes in FMRP-deficient mice by deletion of the *Cpeb1* gene is mediated by restoration of steady state RNA levels and consequent rebalancing of translational homeostasis. Our data establish an essential role of FMRP in codon optimality-dependent mRNA stability as an important factor in FXS.

## Introduction

Fragile X syndrome (FXS) is the most common form of inherited intellectual disability caused by a single gene mutation (1). FXS is caused by a trinucleotide repeat expansion in the *FMR1* locus, which results in transcriptional silencing and loss of its protein product FMRP (2). In FMRP-deficient mice, protein synthesis in the brain is elevated by ∼15-20% (3, 4), indicating that FMRP represses translation. In the mouse brain, FMRP binds mostly to coding regions of approximately 850 to 1000 mRNAs (5–9), and co-sediments with polyribosomes (5). FMRP has been proposed to repress translation by impeding ribosome translocation (5, 10–13). This hypothesis is based on the evidence that ribosomes associated with many of these mRNAs are resistant to puromycin treatment *in vitro*, which causes premature polypeptide release and thus is an indirect measure of ribosome translocation, and that ribosomes transit at faster rates in FMRP knockout (KO) brain extracts compared to WT (5, 10), FMRP also regulates translation directly or indirectly at the level of initiation (14, 15), RNA splicing (13), editing (16, 17), nuclear export (18, 19), and m^6^A modifications (18–20). However, the relationship of these molecular impairments to the etiology of Fragile X Syndrome, if any, is unknown.

Translation is also controlled by the supply and demand of available tRNAs (21). When particular codons represented in mRNA are not met by a sufficient supply of charged tRNA, ribosome transit slows or stalls, which in turn causes RNA destruction (22–27). Control of specific translation and RNA destruction by codon bias varies with tissue (24), time of development (23), and cell stress (28). Additionally, in yeast, trans-acting factors can influence codon bias-mediated RNA destruction (22), indicating a more complex regulation than simple codon-tRNA balance.

Here, we have used ribosome profiling and RNA-seq to investigate translational dysregulation in the FMRP KO cortex and found that FMRP coordinates the link between RNA destruction and codon usage bias (codon optimality). We find that the apparent dysregulation of translational activity (i.e., ribosome occupancy) in FMRP KO cortex can be accounted for by commensurate changes in steady state RNA levels. Downregulated mRNAs in FMRP KO cortex are enriched for those that encode factors involved in neuronal and synaptic functions, and are highly enriched for FMRP binding targets. These observations suggest that in the cortex, FMRP directly or indirectly regulates RNA stability. Indeed, RNA metabolic profiling by 5-ethynyl uridine incorporation and whole transcriptome sequencing reveals wide-spread RNA degradation in *Fmr1* KO cortical neurons while synthesis and processing rates remained substantially unchanged. Of the ∼700 mRNAs that degraded significantly faster in FMRP KO cortex compared to WT, those enriched for optimal codons were particularly affected. This widespread codon-dependent RNA destruction in FMRP-deficient neurons involves a massive reshuffling of the identities of stabilizing or destabilizing codons. Moreover, FMRP can distinguish between optimal and non-optimal codon-containing mRNAs, probably through the associated translational machinery. Finally, we demonstrate that in a genetic rescue paradigm of FXS where a double deficiency of FMRP and CPEB1 mitigates the disorder in mice, restoration of RNA levels drives the recovery of ribosome occupancy. These results indicate that a primary consequence of FMRP depletion from the brain is the uncoupling of codon bias from the RNA destruction machinery. This uncoupling may be a general mechanism that underlies FXS, and restoration of the RNA stability landscape could be a key to ameliorating the disorder.

## Results

### Steady state RNA level changes drive translational buffering in Fragile X brain cortex

To identify mRNAs that are translationally dys-regulated in FMRP-deficient mouse cortex, we performed ribosome profiling (29) and RNA-seq from WT and *Fmr1* KO (FK) animals (**Fig 1a**). Ribosome occupancy (RO), defined as ribosome protected fragments (RPFs) normalized to mRNA levels, is a measure of translational activity (29) and has served as a proxy for protein synthesis. Accumulating evidence suggests that one mechanism whereby FMRP inhibits translation is by stalling ribosome transit (5, 10, 12, 13) and indeed there is a moderately (10-15%) higher rate of protein synthesis in FMRP-deficient brain (3, 4, 30). Using Xtail (31), an algorithm that tests for differential ribosome occupancies (DROs) between samples, we identified 431 mRNAs with DROs between FK and WT (FDR < 0.05; **Fig 1b, Fig S1a**). Consistent with FMRP acting as a translation repressor, 80% of these mRNAs (345/431) have increased RO.

**Figure 1:**
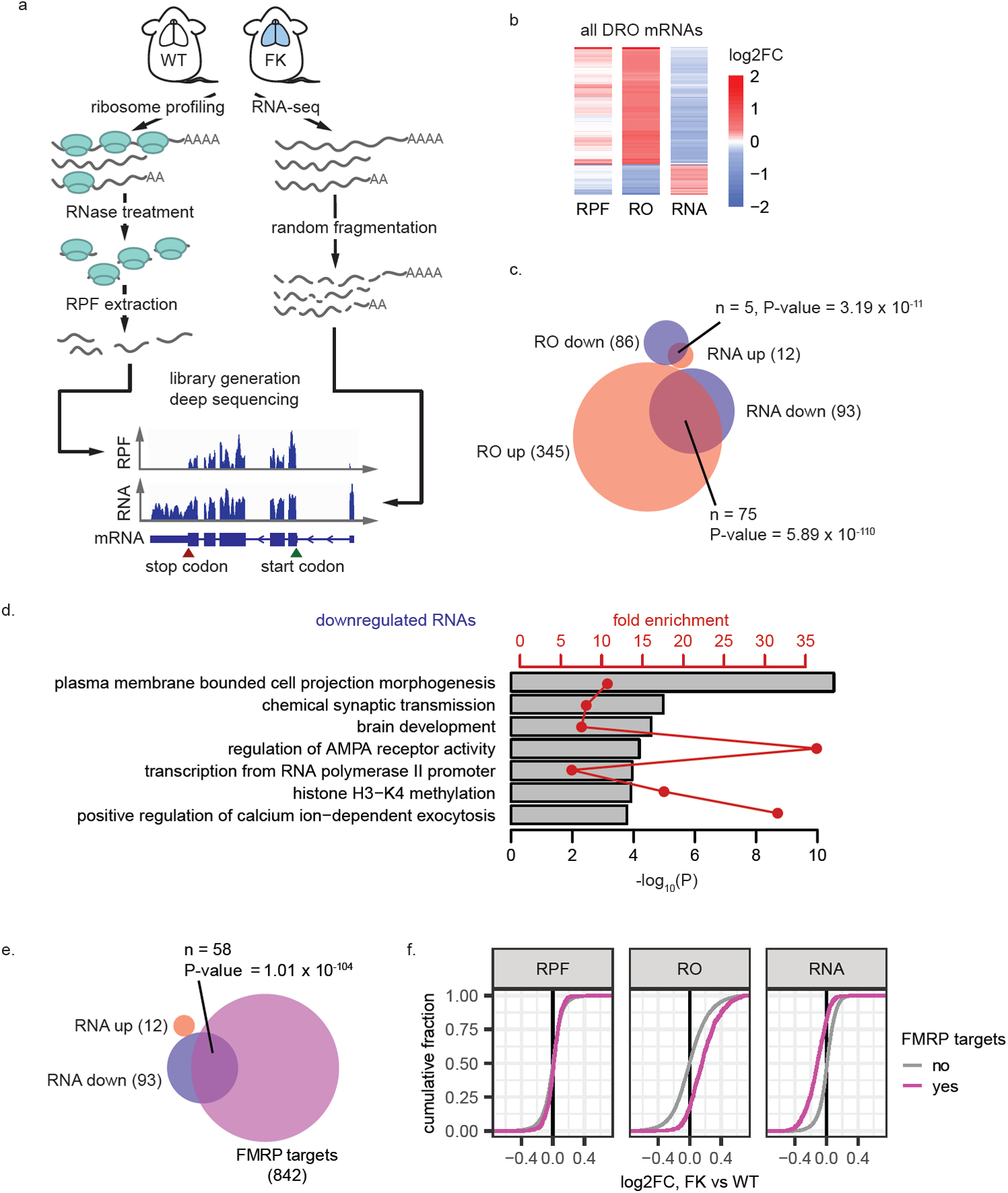
Steady state RNA level changes drive translational buffering in Fragile X brain cortex. **a**, Illustration of the experimental pipeline of ribosome profiling and RNA-seq for WT and FMRP-knockout (FK) mouse brain cortices. RPF: ribosome protected fragments. **b**, heatmaps showing log2FC of RPF, RO (ribosome occupancy), and RNA levels for mRNAs having statistical significant different ROs as calculated using Xtail (31) between FK and WT (2 biological replicates for each genotype, FDR < 0.05). **c**, Venn diagram showing the overlap of mRNAs with DRO and mRNAs that are differentially expressed. **d**, Representative Gene Ontology (GO) terms enriched for mRNAs down regulated at the RNA level in the FK brain cortex. Grey bars and red point-and-lines show the –log_10_(P value) and fold enrichments of each of these GO terms, respectively. See **Table S1** for full lists of enriched GO terms. **e**, Venn diagrams showing the overlap between the downregulated mRNAs and FMRP binding targets (5). Numbers of RNAs in each group and in each overlap as well as p-values of enrichment (hypergeometric test, upper tail) are indicated. **f**, ECDF plots showing log2FC of RPF, RO, and RNA levels for FMRP binding targets (5) and other mRNAs.

To determine the underlying cause of DRO, we analyzed our RPF and RNA-seq data separately. DRO can result from translational dysregulation driven by differential RPF but with little change in RNA levels; conversely, translational buffering occurs when RPFs are unchanged but the DRO is driven by dysregulated RNA levels (32). The RPF changes in the DRO mRNAs are subtle, however, the changes in RNA levels strongly oppose the changes in RO (**Fig 1b**). When taken as a group, these mRNAs with altered ROs have stronger changes at RNA levels than at RPF levels (**Fig S1a**).

We identified 12 and 21 mRNAs with strongest increase and decrease at RPF level, and 12 and 93 mRNAs with strongest increase and decrease at steady state RNA level (including *Fmr1* mRNA) (see **Material and Methods**). The 345 RNAs with increased RO in FK are significantly enriched for those with increased RPFs (n = 7, p-value = 4.17 × 10^−11^, hypergeometric test), but much more so for mRNAs with reduced steady state levels (n = 75, p-value = 5.89 × 10^−110^, hypergeometric test). This is also the case for mRNAs with decreased RO (**Fig 1c**). These data show that mRNA steady state level changes drive the observed RO changes, likely via translational buffering in the FK brain cortex.

Some of the top mRNAs downregulated at the steady state level include those that are involved in ion channel function (e.g. Pdzd2 (33) and Wnk2 (34)) and synapse development and communication (e.g. Bai2 (35) and Sipa1l3 (36)). Gene Ontology (GO) analysis of downregulated mRNAs show an enrichment for functions related to neuronal dendrites, pre- and post-synapses, and channels and receptors (**Fig 1d; Table S1**). The loss of these RNAs could contribute to the neurological defects related to FXS. The FMRP binding target RNAs in the brain are also highly enriched for these similar GO terms (5). Indeed, over two-thirds of the downregulated mRNAs (58/93) are FMRP targets (5) (**Fig 1e**, enrichment p-val = 7.53 × 10^−51^, hypergeometric test), including the four example genes mentioned above. As a group, the FMRP target RNAs are significantly reduced (mean log2FC = −0.12, p-val < 2.2 × 10^−16^, one-tail t test). This observation indicates that the increased RO is caused by reduced RNA levels (mean log2FC = 0.17, p-val < 2.2 × 10^−16^, one tail t test), although their changes at RPF level is very subtle (mean log2FC = 0.0093, p-val = 0.00292, one-tail t test) (**Fig 1e**). These data show that in the FMRP-deficient mouse brain cortex, translational buffering is driven by steady state mRNA changes, particularly the downregulation of the FMRP target mRNAs. Interestingly, the downregulation of mRNAs, including FMRP binding targets, is observed across multiple FXS mouse and human models (**Fig S1**), suggesting that reduction of these mRNAs could underlie FXS.

### RNA metabolic profiling reveals disrupted RNA stability in FMRP-deficient neurons

Because FMRP is mostly a cytoplasmic protein (11), we hypothesized that the downregulation of its target (and other mRNAs) is due to a post-transcriptional mechanism, possibly destabilization upon loss of FMRP. To test this hypothesis, we incubated WT and FK mouse cortical neurons (14 DIV) with 5-ethynyl uridine (5EU) for 0 (i.e., unlabeled control, or “unlab”), 20 (library A), or 60 min (library B), after which the RNA was “clicked” to biotin and purified by streptavidin chromatography. The RNA was mixed with 5EU-labeled *Drosophila* RNA and unlabeled yeast RNA as controls and sequenced together with total unenriched RNAs as input samples (**Fig 2a**). The spike-in *Drosophila* and yeast RNAs for library generation were used as quality control measures, showing that the WT and FK libraries were of equal quality (**Fig S2a-c**). After filtering (**Fig S2d**), we calculated RNA metabolism rates (synthesis, processing, and degradation) by comparing nascent and mature RNA concentrations in the 5EU-labeled and input total RNA libraries using the INSPEcT algorithm (37, 38). We obtained metabolism rate information for 8590 RNAs, which include 412 FMRP target mRNAs. The rates follow log-normal distributions with medians of 1.12 and 1.04 RPKM/hr for synthesis, 6.84 and 6.60 hr^-1^ for processing, and 0.13 and 0.14 hr^-1^ for degradation for libraries A and B, respectively (**Fig S2e**). These values demonstrate the reproducibility of the assay.

**Figure 2:**
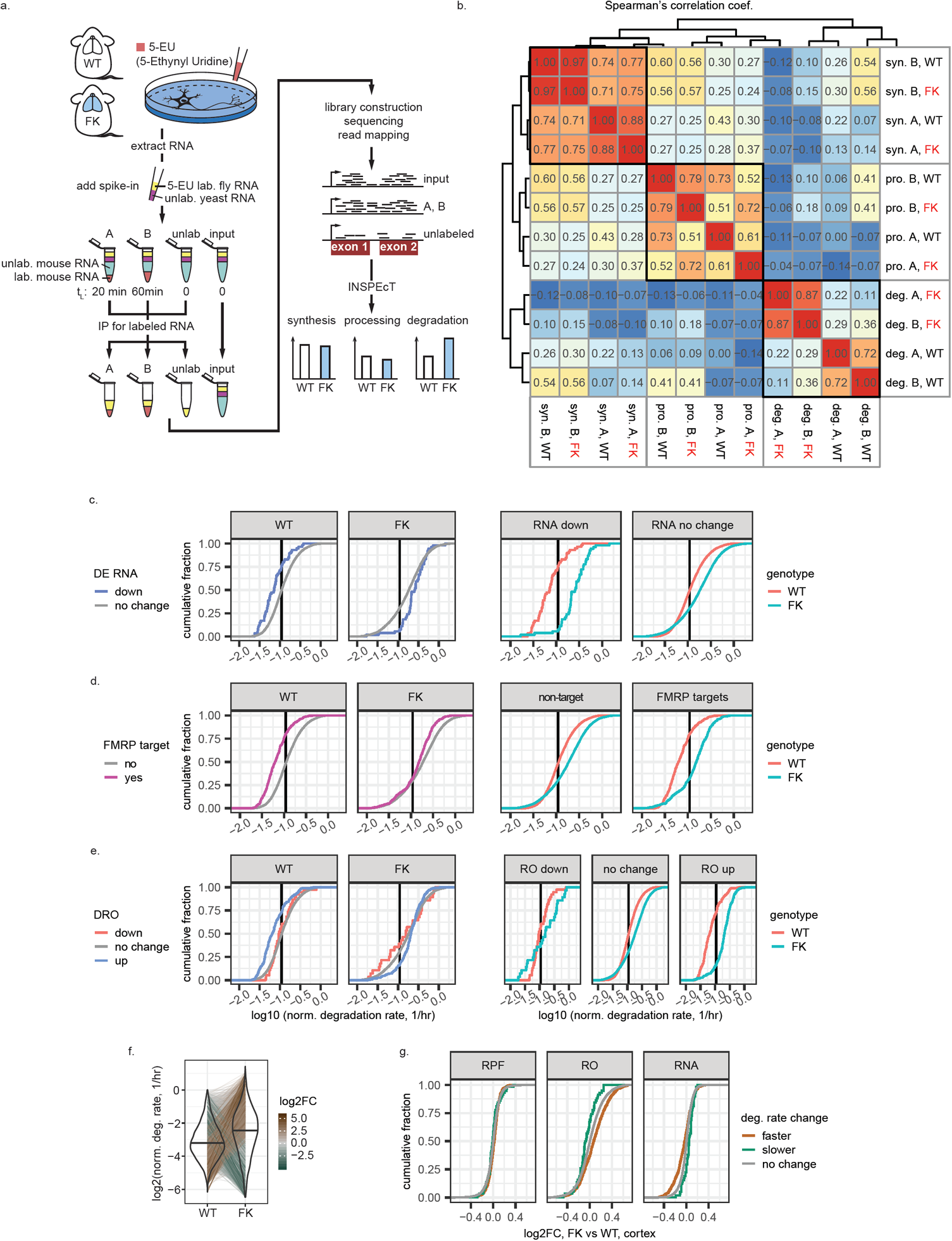
Increased mRNA degradation rates in FK neurons correlate with the steady state RNA depletion. **a**, Illustration of the experimental pipeline of RNA metabolism profiling for WT and FK neurons. Unlab./lab.: unlabeled/labeled. tL: time labeled. **b**, Heatmap of Spearman’s correlation coefficients between synthesis (syn.), processing (pro.), and degradation (deg.) rates estimated from RNA-seq libraries generated from WT and FK neurons labeled for 20 (A) or 60 (B) minutes. Dendrogram shows the unsupervised hierarchical clustering using their Spearman’s correlation coefficients. **c-e**, ECDF plots of normalized degradation rates for downregulated mRNAs in FK cortex (**c**), FMRP binding targets (**d**), and mRNAs with DROs (**e**) and other mRNAs, compared between the mRNA groups (left), or between the genotypes (right). The mean of the normalized degradation rates from libraries A and B of each mRNA was used. Black vertical lines denote the median degradation rates for all mRNAs. **f**, Violin-and-line plot for the degradation rates of all mRNAs. The mean of the normalized degradation rates from libraries A and B of each mRNA was used. The black horizontal line in each violin denotes the median. Thin lines span WT and FK connect the values of the same mRNA in both genotypes. Brown-grey-green shades of the thin lines indicate the log2FC of the normalized degradation rates of each mRNA. **g**, ECDF plots showing log2FC of RPF, RO, and RNA levels in FK cortex compared to WT for mRNAs with faster (brown) or slower (green) degradation rates.

We calculated Spearman’s correlation coefficients for all three metabolism rates for the two genotypes for both libraries (**Fig 2b**). For synthesis, processing, and degradation, we observed decreasing correlation coefficients between WT and FK. For synthesis rates, WT and FK cluster together for the same labeling parameter (library A or B), indicating that there is little genotype difference. For libraries A and B, the correlation coefficients were 0.97 and 0.88 between WT and FK, again demonstrating that the synthesis rates between the two genotypes are similar. For processing rates, the two genotypes were also similar despite slightly lower Spearman’s correlation coefficients between WT and FK (0.79 and 0.61 for libraries A and B). Strikingly, the correlation coefficients for degradation rates between WT and FK were substantially lower (0.22 and 0.36 for libraries A and B), indicating that there is a major difference in RNA degradation between genotypes. The Spearman’s correlation coefficients between libraries A and B for each genotype (0.87 and 0.72, respectively) indicate high reproducibility. Therefore, the degradation rates for the four libraries are separated by genotype (**Fig 2b**), demonstrating that RNA stability in FK neurons is disrupted.

We determined whether the following groups of mRNAs have altered degradation rates: 1) mRNAs that are downregulated (**Fig 2c**), 2) FMRP targets (**Fig 2d**), and 3) mRNAs that have DROs (**Fig 2e**). For convenience, we will refer to group 1 as downregulated mRNAs and group 3 as RO up/down mRNAs, respectively. Statistical tests were performed on log10 transformed degradation rates. In WT neurons, the downregulated mRNAs degraded significantly slower than the other RNAs (p-val = 1.31 × 10-6, two-tailed t test). However in FK neurons, the downregulated mRNAs degraded significantly faster (p-val = 0.000684, two-tailed t test) (**Fig 2c, left**). Moreover, the mRNAs in FK neurons are globally destabilized relative to those in WT neurons (for the mRNAs not downregulated in FK cortex, the difference of log10 degradation rate means in FK vs WT = 0.17 1/hr, p-value < 2.2 × 10^−16^, two-tailed t test), however the downregulated mRNAs are destabilized even more (difference of log10 degradation rate means in FK vs WT = 0.53 1/hr, p-value = 2.09 × 10^−15^, two-tailed t test) (**Fig 2c, right**). FMRP targets and RO up mRNAs are also destabilized more than the global trend (**Fig 2d, e**). Interestingly, the RO down mRNAs were resistant to mRNA destabilization upon loss of FMRP; their degradation rates are not significantly different in FK compared to WT neurons (p-val = 0.323, two-tailed t test) (**Fig 2e, right**). This resistance to transcriptome-wide destabilization could explain the observed steady state upregulation of these mRNAs (**Fig 1b**).

We did not identify RNAs with dysregulated synthesis or processing rates in FMRP-deficient neurons, but detected 748 RNAs with altered degradation rates, of which 688 (92%) degraded faster in FK compared to WT (adjusted p-value cut-off of 0.01) (**Fig 2f**). The RNAs that degraded faster in FK neurons were significantly enriched for FMRP targets as well as those that were downregulated at the steady state level in the cortex (**Fig S2g**). As groups, the RNAs that degraded faster or slower in FK neurons displayed decreased or increased steady state levels (p-val = 5.49 × 10^−14^ and 1.47 × 10^−4^, two-tailed t test), as well as increased or decreased ROs in FK cortex, respectively (p-val < 2.2 × 10^−16^ and = 5.28 × 10^−3^), with no change at the RPF level (**Fig 2g**). Based on these data, we conclude that there is widespread mRNA destabilization, including many of the FMRP targets, in the absence of FMRP; this in turn drives steady state RNA level changes and translation buffering.

### Loss of FMRP uncouples the link between optimal codons and mRNA stability

To identify features of mRNAs that are involved in this FMRP-dependent stability, we examined the correlation of brain cortex steady state mRNA level changes or neuronal mRNA degradation rate changes with codon optimality (gene codon Adaptation Index (cAI) score, see **Material and Methods**), coding sequence (CDS) guanine-cytosine (GC) content, CDS, 5’ and 3’ untranslated region (UTR) lengths and their minimal energy of folding (MEF, an indication of possible secondary structure) (**Fig S3a**). The strongest and most consistent correlations with both cortical steady state mRNA level changes and neuronal mRNA degradation rate changes were gene cAI scores and CDS GC content, which are correlated features themselves (39, 40). More specifically, the log2FC of cortical steady state mRNA levels had a negative correlation with gene cAI scores (Spearman’s correlation coefficient = −0.22, p-value < 2.2 × 10^−16^) while the log2FC of the neuronal mRNA degradation rate had a positive correlation (Spearman’s correlation coefficient = 0.20, p-value < 2.2 × 10^−16^) (**Fig S3a**). This means mRNAs which contain more optimal codons were the ones that had faster degradation rates and showed a consequent downregulation in FK.

Codon optimality, a measure of the balance between the usage frequency of a given codon (demand) and supply of charged tRNAs encoding the complementary anticodon (21), is a major determinant of mRNA stability from yeast to vertebrates (22–27). Generally, mRNAs with more optimal codons (high gene cAI scores; presumably with faster decoding rates) are more stable than mRNAs using less optimal codons, connecting translation regulation to mRNA stability. Because FMRP regulates translation, codon optimality could be a mechanism that links FMRP-mediated translation to mRNA stability. Therefore, we investigated the role of codon optimality in FMRP-dependent mRNA stability.

We grouped all detectable mRNAs into 10 equal sized bins of increasing gene cAI score and examined which bins were particularly affected by mRNA degradation rates (**Fig 3a**). mRNAs in bins 4-10 degraded significantly faster in FK neurons than in WT neurons (Holm adjusted p-value < 0.01, one tailed-t test); the higher the cAI score bin, the faster the rate of degradation. However, mRNAs in bins 1-3 were barely changed, indicating that mRNAs containing non-optimal codons are resistant to destabilization upon loss of FMRP.

**Figure 3:**
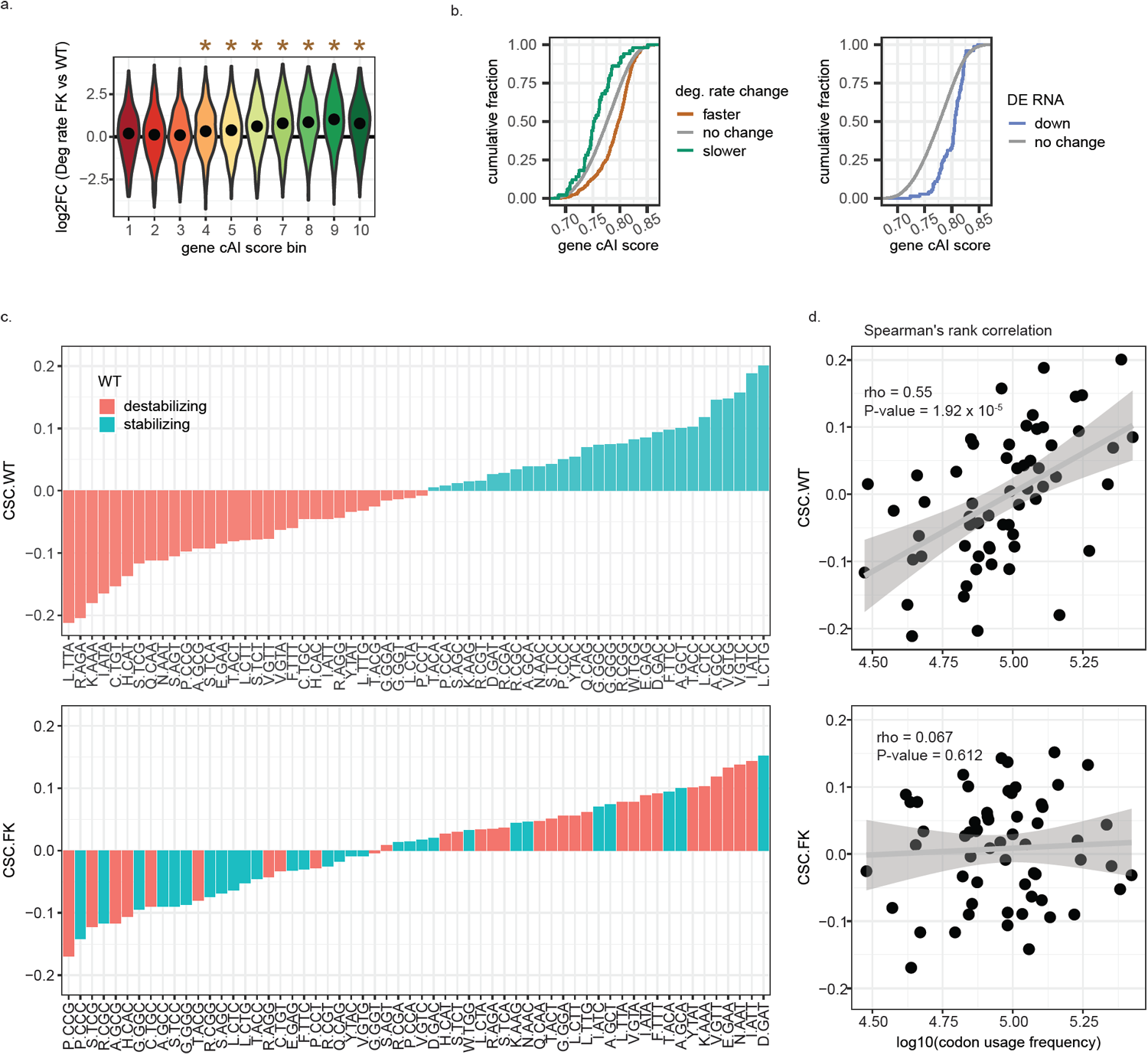
Increased mRNA degradation rates in FK neurons are codon optimality dependent. **a**,, All mRNAs were grouped into 10 equal bins based on their gene cAI scores. Bin 1 contains genes with gene cAI scores of the lowest quantile and bin 10 contains genes of the highest quantile. Violin plots show log2FC of degradation rates in FK vs WT neurons for mRNAs in each gene cAI score bins. The point in each violin denotes the median of the bin. Brown stars indicate the median of the bin greater than 0 (Holm adjusted p-value < 0.01, one tailed-t test). No bin had median less than 0. **b**, ECDF plots of gene cAI scores for mRNAs with faster or slower degradation rates in FK neurons (left) or reduced mRNA levels in FK cortex (right). **c**, Bar graphs for Codon-Stability Coefficients (CSC) for each codon as arranged from minimum to maximum in WT (upper) and FK (lower) neurons. The color of each bar indicates the codon as stabilizing (CSC > 0, green) or destabilizing (CSC < 0, orange) in WT neurons. The amino acid for each codon is indicated. **d**, Scatter plots and linear regressions of the CSCs as a function of log_10_(codon usage frequency) of the top 10% of expressed genes in WT (upper) and FK (lower) neurons. Spearman’s rank correlation coefficients and p-values of the correlations are indicated.

Compared to the general transcriptome (gene cAI scores = 0.78 ± 0.038, mean ± standard deviation), mRNAs that degrade faster in FK neurons (gene cAI scores = 0.79 ± 0.033) and the mRNAs that are downregulated in the cortex (gene cAI scores = 0.80 ± 0.023) have significantly more optimal codons (p-val < 2.2 × 10-16 and 1.12 × 10-14, two-tailed t test). However, the mRNAs that degrade slower in FK neurons (gene cAI scores = 0.76 ± 0.034) are significantly less optimal (p-val = 2.85 × 10-4, two-tailed t test) (**Fig 3b**). These results indicate that FMRP stabilizes mRNAs that have an optimal codon bias.

Because optimal codons may confer stability to mRNAs (22–27), we examined whether this link is uncoupled upon loss of FMRP. The codon-stability coefficient (CSC), which describes the link between mRNA stability and codon occurrence, has been calculated for each codon from yeast to human (23, 24, 26, 27). We determined whether this relationship is maintained in the absence of FMRP. The CSC values in WT neurons ranged from < −0.2 to > 0.2, which is comparable to previously reported CSC values for human cell lines and mouse embryonic stem cells (**Fig S3b**). This differs from what has been described in *Drosophila* where the neuronal CSC is attenuated relative to somatic cells (24). Of the 60 non-start or -stop codons, 29 had CSCs greater than 0 (stabilizing codons) and 31 had CSCs smaller than 0 (destabilizing codons) (**Fig 3c**, upper). Strikingly, 17 codons that are stabilizing in WT are destabilizing in FK neurons, and 21 codons changed in the opposite direction (**Fig 3c**, lower). Optimal codons are prevalent in abundant mRNAs and are associated with positive CSCs (i.e., are stabilizing codons) (23, 24, 26, 27), as we have observed in WT neurons (**Fig 3d upper**) (Spearman’s r = 0.55, p-value = 1.92 × 10^−5^). In FK neurons, however, this relationship breaks down. Codon usage frequencies in the top 10% of highly expressed mRNAs are the same as in WT (data not shown), but the correlation between codon usage frequencies and CSCs is nearly random in FK (r = 0.067, p-value = 0.612) (**Fig 3d**). These results show that the link between codon-mediated RNA stabilization and codon usage bias is uncoupled in FMRP-deficient neurons.

### FMRP may stabilize codon optimality dependent mRNA stability directly and through other factors

We considered possible mechanisms for how FMRP could regulate codon optimality-dependent mRNA stability. Because the downregulated mRNAs are enriched for FMRP targets, FMRP may stabilize them through direct binding. We noticed a bias for high gene cAI scores of the FMRP targets (**Fig 4a**) and hypothesized that FMRP may associate with mRNAs of higher codon optimality. To test this, we compared a pair of previously described reporter mRNAs that are enriched in optimal or non-optimal codons (27). They differ by a single nucleotide insertion to induce a frame-shift, and thus they have almost identical nucleotide compositions. HEK293T cells were transduced with these reporters together with a FLAG-tagged FMRP-expressing vector (41). Subsequent anti-FLAG immunoprecipitation and RT-qPCR showed that the reporter with optimal codons was more enriched compared to the reporter with non-optimal codons (p-value = 0.0261, one-tailed paired t test; **Fig 4b**). This result suggests that FMRP can indeed “identify” mRNAs with optimal codons as opposed to general nucleotide composition.

**Figure 4:**
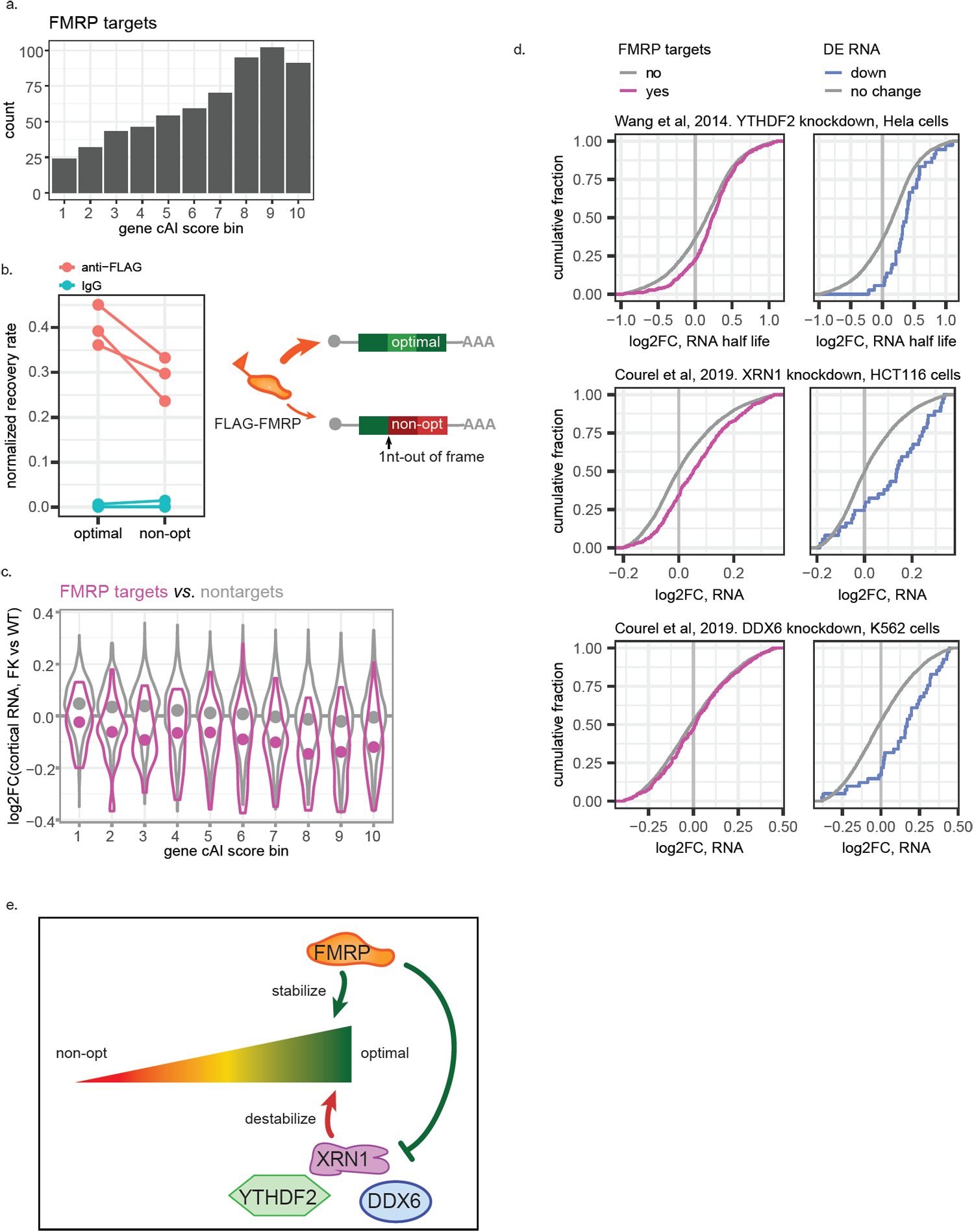
FMRP and other factors may regulate optimal codon dependent mRNA stability. **a**, Bar graph of count of FMRP target genes in each gene cAI score bin. **b**, RNA immunoprecipitation (RIP) testing FMRP recruitment to reporter mRNAs that use optimal versus nonoptimal codons. The reporter that uses nonoptimal codons was generated using 1nt insertion into the reporter that uses optimal codon (left) (27). Normalized recovery rate was calculated by normalizing the recovery rate of the reporter by the recovery rate of the *Fmr1* mRNA, the latter as an internal control for the RIP experiment. Each point, a biological replicate, is an average of 2-3 technical replicates. Each line connects data from the two reporters in a paired experiment that was performed independently from other pairs. p-value = 0.0261, one-tailed paired t test. **c**, Violin plots show log2FC of steady state RNA levels in FK vs WT cortex for all FMRP targets (purple) or non-targets (grey) in each gene cAI score bins. The point in each violin denotes the median of the bin. **d**, ECDF plots showing log2FC of RNA levels for FMRP binding targets (left) and mRNAs downregulated in FK cortex (right) after knock-down of YHDF2 (42), XRN1, and DDX6 (43) in human cell lines. Only genes with unique mouse orthologs were considered. **e**, model of how FMRP could stabilize mRNAs that use optimal codons directly and through other factors.

We next examined whether FMRP binding could impact RNA stability. As a group, the degradation rate of FMRP CLIP targets was higher than the non-target RNAs (p-value = 1.36 × 10^−9^, one-tailed t test; **Fig S4a**). Because too few FMRP targets (30%) were detected in the degradation rate data set, we focused on the steady state RNA changes and compared RNAs in each cAI score bin. Similar to the global trend discussed above (**Fig 3a**), FMRP targets with more optimal codons were also more strongly downregulated than those with non-optimal codons (Spearman’s correlation coefficient of RNA log2FC and gene cAI score bin = −0.21, p-value = 8.27 × 10^−8^). However, FMRP targets in all but the lowest codon optimality bin were significantly reduced in the absence of FMRP (Holm adjusted p-value < 0.01, one-tailed t test) (**Fig 4c, red**). This is different from the codon optimality matched non-targets (**Fig 4c, black**). The non-target mRNAs also showed an optimal codon biased downregulation (Spearman’s correlation coefficient of RNA log2FC and gene cAI score bin = −0.20, p-value < 2.2 × 10^−16^), but only RNAs in cAI bins 7, 8, and 9 are significantly downregulated (Holm adjusted p-value < 0.01, one-tailed t test). In all gene cAI score bins, the log2FC of the FMRP targets are significantly more negative than the non-target RNAs (Holm adjusted p-value < 0.01, one-tailed t test). These results suggest that the stability of the FMRP target RNAs is not only affected by their codon optimality, but also by the FMRP binding.

Not all mRNAs that are depleted in the FK cortex are FMRP CLIP targets, suggesting that FMRP could also stabilize mRNAs through other factors. For example, YTHDF2 was shown to antagonize FMRP’s stabilizing effect for some of FMRP targets in mouse brain cortex (20), and knockdown of this protein in human cell lines (42) leads to longer half-lives of the FMRP targets (D = 0.14, p-value = 2.19 × 10^−7^, two-sided KS test) and of mRNAs downregulated in the FK cortex (D = 0.37, p-value = 3.56 × 10^−5^, two-sided KS test) (**Fig 4d**).

We examined RNA-seq datasets to assess other factors that might be involved in FMRP-mediated optimal codon-dependent mRNA stability. We selected factors that target mRNAs of higher GC content (43) or higher codon optimality (44), and therefore the RNA level changes upon depletion of these factors likely affects their stability (**Fig 4d, Fig S4b**). Depletion of XRN1 (5’−3’ exonuclease) and DDX6 (DEAD-Box Helicase 6) in human cell lines (43) leads to upregulation of the mRNAs downregulated in FK cortex (D = 0.42, p-value = 4.05 × 10^−8^; D = 0.37, p-value = 2.24 × 10^−6^, two-sided KS test) (**Fig 4d**). On the other hand, depletion of FXR1 (Fragile X Mental Retardation Syndrome-Related Protein 1) (45) leads to downregulation of the orthologs of FMRP targets (**Fig S4b**). These results suggest that XRN1 and DDX6 might antagonize FMRP while FXR1 might cooperate with it to stabilize mRNAs of high codon optimality.

### CPEB1 depletion rescues FXS by rebalancing RNA homeostasis

Nearly all pathophysiologies associated with FMRP deficiency are rescued in FMRP/CPEB1 double knockout (dKO) mice (10). To assess how this rescue could be achieved, we performed ribosome profiling and RNA-seq on CPEB1 KO (CK) and dKO mouse brain cortices in addition to the WT and FK samples as described above (**Fig 5a**). We identified 651 genes with DROs among the four genotypes (FDR < 0.05; **Fig 5b**), including the 431 genes with DRO between FK and WT described above (**Fig 1b**). Importantly, 425 of these DROs were rescued in the dKO cortex (i.e., not significantly different compared to WT). Unexpectedly, >50% of RNAs with DRO in CK (204 out of 359) also had DROs in FK, and were changed in the same direction (i.e., increased or decreased). These molecular data are consistent with previous observations such as dendritic spine number and metabotropic glutamate receptor-mediated long term depression (mGLurRLTD), which are nearly identically aberrant in the two single KO mice but rescued to normal in dKO animals (10). Because of the molecular similarities between WT and dKO, and between FK and CK, we henceforth refer to these two groups as “normal” and “FXS-like.”

**Figure 5:**
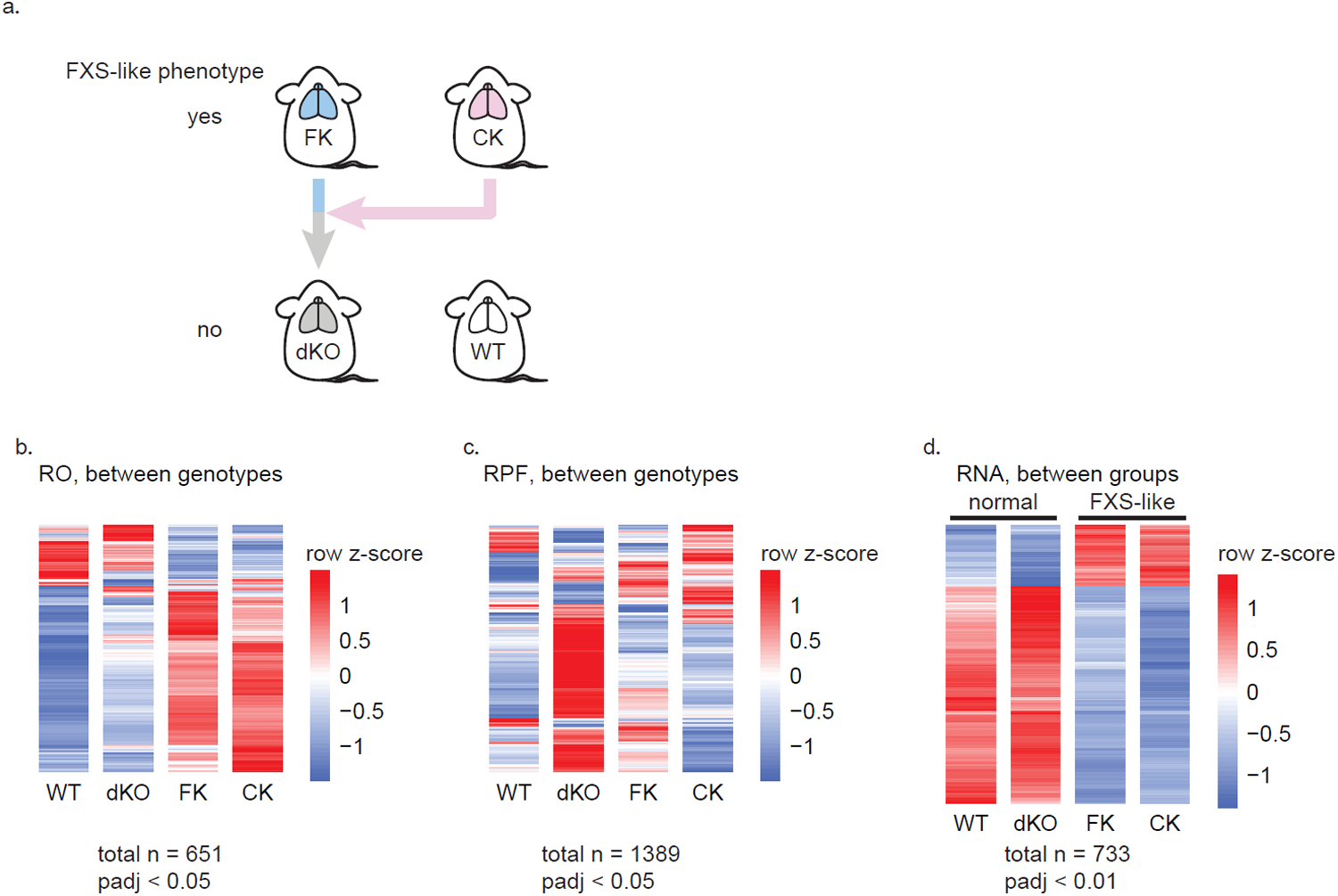
Genetic rescue of FX by CPEB KO is rebalanced at RNA level. **a**, illustration of the genetic rescue paradigm described previously (10). CK, CPEB1 KO; dKO, double KO. **b-c**, Heatmaps showing mRNAs having differential RO (**b**) and RPF (**c**) between any two genotypes of the four genotypes noted above. **d**, Heatmaps showing mRNAs that are differentially expressed between the normal (WT and dKO) and FXS-like groups (FK and CK). Red and blue shades of the heatmaps show high or low z-scores calculated for each mRNA (row) across all samples. Each genotype has 2 biological replicates.

To determine the underlying molecular driver of the DRO among the four genotypes, we analyzed our RPF and RNA-seq data separately. At padj < 0.05, only 23 and 21 RPFs were significantly different between FK and WT and between CK and WT. Conversely, the dKO was the most different from WT with 410 and 333 RPFs that were significantly higher or lower (**Fig 5c**). Therefore, this genetic rescue of FXS-like phenotypes in the dKO is not achieved by correcting translational activity as defined by the number of RPFs. In contrast, the RNA-seq heatmap displayed a mirror image of the DRO heatmap (**Fig S5a)**. Compared to WT, the expression of 50 genes was dys-regulated in FK (padj < 0.05; 10 up-regulated, 40 down-regulated), 145 in CK (padj < 0.05; 13 upregulated, 132 downregulated), but only 2 in dKO (padj < 0.05; *Cpeb1* and *Fmr1*). The differentially expressed (DE) genes in FK and CK were largely identical. Among the 10 and 13 genes up-regulated in FK and CK, 7 overlap (p = 8.72 × 10^−25^, hypergeometric test, upper tail); among the 40 and 132 genes down-regulated, 35 overlap (p = 3.91 × 10^−72^, hypergeometric test, upper tail). Because the transcriptome profiles in FK and CK are as similar as the WT and dKO profiles, we performed an unsupervised hierarchical clustering to test for sample to sample similarities (**Fig S5b)**. FK and CK formed one cluster while WT and dKO formed another, validating the “FXS-like” vs “normal” grouping at the RNA level.

Having validated the grouping, we tested for DE RNAs in the FXS-like group (FK and CK) relative to the normal group (WT and dKO). The DE RNAs between the groups are changed the same direction (i.e., up or down) in the single KOs and are rescued in the dKO to WT levels. With greater statistical power than when comparing between single genotypes, we identified 733 dys-regulated RNAs in the FXS-like group (padj < 0.01), 162 (22.1%) up-regulated and 571 (77.9%) down-regulated (**Fig 5d)**. Almost all downregulated RNAs in FK compared to WT are downregulated in the FK-like group (87/93). Over 77% of the RNAs with up-regulated ROs in FK vs WT (267 out of 345) were significantly reduced in the FXS-like group (p-value = 0, hypergeometric test, upper tail). Similarly, 42% of the RNAs with down-regulated ROs in FK (36 out of 86) were significantly increased steady state levels (p-value = 3.28 × 10^−51^, hypergeometric test, upper tail). Therefore, we conclude that rescue of FXS in FMRP/CPEB1 dKO animals is achieved by rebalancing the transcriptome.

## Discussion

It is widely assumed that excessive protein synthesis is not only a corollary of FXS, but a proximate cause of the disorder (46). Consequently, several studies have used high resolution whole-transcriptome methods to identify mRNAs that are either bound by FMRP (5–9) or whose translation is dysregulated in FMRP-deficient mouse models (5, 12, 13, 47, 48). Although nearly 1000 FMRP binding targets have been identified in the mouse brain, surprisingly few are dysregulated at the translational level, and of those that are, the dysregulation is usually modest. Moreover, in even fewer cases has the dysregulation been linked to FXS pathophysiology (49). Therefore, the functional output of FMRP binding remains largely elusive. Our study shows that in the FK mouse brain cortex, steady state mRNA levels are globally disrupted, which drive the dysregulated translational buffering (**Fig 1, 5**). FMRP targets are particularly downregulated, which happens specifically in neurons (50). By metabolic labeling and RNA-seq in neurons, we show that the loss of FMRP results in reduced stability not only of its direct target substrates, but also of other mRNAs with an optimal codon bias (**Fig 2, 3**). This instability drives the changes in steady state RNA levels and translational buffering in the cortex (**Fig 2**). Moreover, our data demonstrate that RNA stability conferred by optimal codons is mediated by FMRP and possibly other trans-acting factors. The loss of FMRP leads to a massive reshuffling of the identities of stabilizing versus destabilizing codons (**Fig 3**).

Rebalancing of RNA levels is accompanied by and probably necessary for mitigation of FXS-like phenotypes in mice by CPEB1 depletion (**Fig 5**) (10). Among the mRNAs dysregulated between the FXS-like group (i.e., FK and CK) and the normal group (WT and FK/CK dKO), GO analysis shows that many up-regulated and rescued mRNAs have protein synthetic functions including ribosome biogenesis, translation, and protein folding, while the down-regulated and rescued mRNAs have cell projection, synaptic transmission, as well as transcription and chromatin functions (**Fig S6a; Tables S2-3)**. Several important points come from this analysis. First, the up-regulated mRNAs are among the highest expressed in the brain (**Fig S6b**), and their protein products could promote general protein synthesis. This could explain the net increase in protein output in the FXS brain (3, 4, 30). Second, we find that FMRP regulates the levels of mRNAs that encode chromatin modifying factors, which is reminiscent of other observations showing that FMRP controls the synthesis of epigenetic regulators, albeit at the translational level (13, 49). Third, the brain and neuron related GO terms enriched for the down-regulated RNAs reflect the neural dysfunction that occurs in FXS. Indeed, the down-regulated RNAs are also significantly enriched for those related to autism as compiled by the Simons Foundation SFARI project (51) (**Fig S6c**; p = 3.56 × 10^−7^, hypergeometric test, upper tail). Therefore, restoring RNA homeostasis could be a key to FXS treatment.

How does FMRP stabilize mRNAs that use optimal codons? One possibility is that FMRP can stabilize its targets by directly interacting with the translational machinery. We show that FMRP preferentially binds an mRNA reporter that contains optimal codons over a reporter that contains non-optimal codons but has nearly the same nucleotide composition (**Fig 4b**). Upon loss of FMRP, target mRNAs are destabilized more than non-targets matched for their codon optimality (**Fig 4c**), suggesting FMRP binding could lead to enhanced stability in addition to that provided by optimal codons. In yeast, the Ccr4-Not complex directly binds the empty E site of the ribosome during ineffective decoding on a non-optimal codon, thereby rendering the mRNA more susceptible to degradation (52). One can postulate that the E site on an optimal codon is more frequently occupied compared to a non-optimal codon, and therefore is less likely to be recognized by the Ccr4-Not complex. Moreover, the N-terminus of *Drosophila* FMRP was also shown to be able to bind directly to ribosomes (53). If these findings are conserved in mammalian systems, FMRP binding to ribosomes may further reduce the chance of Ccr4-Not binding.

We cannot rule out the possibility that FMRP may “sense” other features of mRNA, e.g. GC content of the coding sequence. In mammalian systems, mRNAs that contain many optimal codons inevitably have higher GC content in the coding sequence, which predicts a stronger secondary structure (40). Interestingly, mRNAs that have reduced steady state RNA levels or have increased degradation rates tend to have long CDS, which also predicts more CDS mRNA structures, as suggested by their low MFE (**Fig S3**). In mammalian cells, strong secondary structure in the CDS stabilizes mRNA, possibly by preventing strand-specific endonucleolytic cleavage or ribosome collisions (40). FMRP binding may help stabilize such structures in the CDS, thereby reducing RNA degradation. Indeed, there is at least *in vitro* evidence that RNA secondary structure is recognized by FMRP (1). On the other hand, DDX6 was shown to mediate mRNA decay of GC rich mRNAs, possibly by facilitating unwinding of the secondary structures throughout the transcripts and facilitating XRN1 progression (39). Therefore, it is possible that FMRP stabilizes the mRNAs by antagonizing DDX6 and XRN1. Indeed, the orthologues of the mRNAs downregulated in the mouse FK brain are upregulated upon loss of DDX6 and XRN1 in human cells (**Fig 4d**), suggesting FMRP, DDX6 and XRN1 may share common targets. By interacting with DDX6, FMRP could regulate a much wider pool of mRNAs than those in the FMRP target list (5).

## Material and Methods

### Animals

WT, FK, CK and dKO mice were as used previously (10). Specifically, FK (JAX stock# 004624) and its WT controls (JAX stock# 004828) were purchased from the Jackson Lab. CK were created in-lab (54). Mice were bred as previously described (10). All mice were maintained in a temperature-(25°C), humidity- (50–60%) and light-controlled (12 hr light-dark cycle) and pathogen-free environment. Animal protocols (#1158) were approved for use by the University of Massachusetts Medical School Institutional Animal Care and Use Committee (IACUC).

### Ribosome profiling and RNA-seq in cortex

Two mice per genotype were used for ribosome profiling and RNA-seq. The brain was rapidly removed from P28–P35 mice, rinsed in ice-cold dissection buffer (1× HBSS + 10 mM HEPES-KOH), rapidly dissected in dissection buffer ice-liquid mixture to collect cerebral cortex as described previously(55). Both cortex hemispheres were homogenized in 900µl of homogenization buffer(55) (10 mm HEPES-KOH, pH 7.4, 150 mm KCl, 5 mm MgCl_2_, 0.5 mm DTT, 100 µg/ml cycloheximide and 2 µg/ml harringtonine), containing protease and phosphatase inhibitors (cOmplete, EDTA-free Protease Inhibitor Cocktail and PhosSTOP from Roche/Sigma, cat. no. 11836170001 and 4906837001), in a pre-chilled 2-mL glass Dounce homogenizer, 20 strokes loose, 20 strokes tight, and centrifuged at low speed (2000 rcf 10 min 4°C) to pellet insoluble material(10). Five hundred microliters of the resulted ∼700µl supernatant (cytoplasmic lysate) were used for ribosome profiling, the rest for RNA-seq. For ribosome profiling, the lysate was digested by 60 U RNase T1 (Thermo Scientific, cat. no. EN0541) and 100ng RNase A (Ambion, cat. no. AM2270) per A260 unit(56) for 30 min at 25°C with gentle mixing. Digestion was stopped by adding 30 µl SUPERase·In (Invitrogen, cat. no. AM2694). Digested lysate was separated by sedimentation through sucrose gradients. Monosome fractions were identified, pooled, and extracted with TRIzol LS (Invitrogen, cat. no. 10296028).

For RNA-seq, cytoplasmic RNA was extracted from the lysate using TRIzol LS. Ten micrograms of RNA were depleted of rRNA using Ribo-Zero Gold rRNA Removal Kit, Human/Mouse/Rat (Illumina, discontinued), and fragmented by incubating with PNK buffer (NEB, cat. no. M0201S) for 10 min at 94°C. Fragmented RNA as separated on 15% urea-polyacrylamide gel, and 50-60nt fraction was collected.

Ribosome profiling and RNA-seq libraries were prepared following published protocols(57) and sequenced with Illumina NextSeq.

### Spike-in RNA for RNA metabolism profiling

*D. melanogaster* (fly) Schneider 2 (S2) cells were grown in 12 ml Schneider’s insect medium (Sigma-Aldrich, cat. no. S0146) containing 10% (v/v) of Fetal Bovine Serum (FBS, Sigma-Aldrich, cat. no. F2442) at 28°C until confluent. Cells were incubated with 200 µM 5-EU for 24 hr, and were washed, pelleted and snap frozen in liquid nitrogen. RNA was extracted using TRIzol.

*S. cerevisiae* (yeast) cells were grown in 10 ml YEP medium containing 3% glucose at 30°C until OD_600 nm_ reaches 0.5. Cells were then pelleted and RNA was extracted using hot acidic phenol (58).

### RNA metabolism profiling with cortical neuron cultures

Cortical cell suspension were obtained by dissociating cerebral cortices from E18 embryos using the Papain Dissociation System (Worthington, cat. no. LK003150). One million live cells were plated in 5 ml complete Neurobasal culture medium (Neurobasal™ Medium (Gibco, cat. no. 21103049), 1x B-27 supplement (Gibco, cat. no. 17504044), 1x Antibiotic-Antimycotic (Gibco, cat. no. 15240096), 1x GlutaMAX (Gibco, cat. no. 35050061)) per 60 mm poly-L-lysine treated cell culture dish. Neurons were fed by half-replacing the complete Neurobasal culture medium twice per week. DIV14 neurons were incubated with 200 µM 5-EU (Click-iT™ Nascent RNA Capture Kit, Invitrogen, cat. no. C10365) for 0 (input and “unlab”), 20 (“A”), and 60 (“B”) min. Neurons were then washed with ice cold 1x PBS buffer, and RNA was extracted using TRIzol (Invitrogen, cat. no. 15596018). Five-EU labeled RNA was enriched and RNA-seq library was prepared by adapting the Forrest et al protocol (59). Specifically, mouse neuron RNA was spiked-in with 10% (w/w) 5-EU labeled fly RNA and 10% (w/w) yeast RNA. The mixed RNA was depleted of rRNA using the Ribo-Zero Gold rRNA Removal Kit (Human/Mouse/Rat) and fragmented using NEBNext® Magnesium RNA Fragmentation Module (NEB, cat. no. E6150S) for 5 min. RNA samples for libraries unlab, A and B were biotinylated by Click-iT chemistry and pulled-down using the Click-iT™ Nascent RNA Capture Kit. The pulled-down samples, together with the input sample, were subjected to library construction the same as for ribosome profiling libraries(57). Here, for the pulled-down samples, all reactions were performed directly on beads until after the reverse transcription step. Sequencing library fraction with insert size 50-200nt was collected and sequenced with Illumina NextSeq. For each WT and FK, two independent batches of neurons were prepared, and each batch resulted in one of each input, unlab, A and B libraries (**Table S4**).

### Differential translation and RNA expression analysis

Brain cortex ribosome profiling and RNA-seq reads were processed as previously described^11^, which includes the following steps: 1) reads were separated based on sample barcode sequences; 2) known 3’ adapter sequences and low quality bases were removed with Cutadapt(60) using parameters -O 2 -q 15 -a “TGGAATTCTCGGGTGCCAAGGAGATCGGAAGAGCGGTTCAGCAGGAATGCCGAGACCG”; 3) reads mapped to rRNA and tRNA genes were removed using Bowtie2(61) with parameter -N 1; 3) remaining reads were mapped to the mm10 genome using TopHat2(62); and 4) PCR duplicates were removed based on Unique Molecular Identifier sequences.

Uniquely mapped reads were then used as input to RSEM(63) for quantification of gene expression and either mapped to RefSeq (v69) mouse coding sequences (RPF) or to whole-transcriptome (RNA-seq). Genes were filtered to have a minimum of 10 TPM (transcripts per million) in at least one sample. We used log transformed TPM expression values to correct for batch effects using ComBat (64) (v3.18.0). Corrected values were transformed back to read counts using the expected size of each transcript informed by RSEM. Batch-corrected counts were used to identify differentially translated/expressed genes with DESeq2 (65) (RPF and RNA) or Xtail (31) (ribosome occupancy, RO). To identify mRNAs with significant DROs, a FDR < 0.05 was used as cut off. To identify mRNAs with strongest changes at RPF and RNA levels, a padj < 0.1 was used as cut off.

### GO analysis

GO enrichment analysis was performed using Cytoscape with the ClueGo(66) plug-in (v2.3.3), with genes that are expressed in the mouse cortex as the reference gene set. Specifically, biological function GO terms of levels 6-13 were tested for enrichment at adjusted p-value < 0.05 (Right-sided hypergeometric test). Enriched GO terms that are similar were then fused to a group based on their Kappa score which quantifies percentage of common genes between terms. The leading group terms, which are the terms with highest significance in each group, are presented in **Fig 1d** and **Fig S6a**. All enriched terms are in **Table S1-3**.

### RNA metabolism profiling analysis

Reads generated from the RNA metabolism profiling libraries were processed as for cortical RNA-seq libraries described above, except that a mouse-fly-yeast merged genome (mm10 + dm6 + sacCer3) was used as the reference genome for reads mapping by hisat2. The mapping statistics here were used for quality control and filtering purposes (**Fig S2a-d**). Uniquely mapped reads that are depleted of rRNA, tRNA sequences and PCR duplicates were again mapped to mm10 genome with hisat2. Intron and exon read quantification, and RNA metabolism rates (synthesis, processing and degradation) estimation was performed using INSPEcT (37) (v1.10.0), with the degDuringPulse parameter set to TRUE. One set of libraries, which was of low complexity (**Table S4**), was still used to confirm the global shift of degradation rates in FK neurons. This reproducible global shift allowed us to normalize WT and FK libraries separately for our gene level analysis (**Fig S2f**). Specifically, raw RNA metabolism rates estimated by INSPEcT were normalized between libraries A and B for WT and FK neurons separately using the limma package(67) with the “cyclicloess” method. After normalization, genes with different metabolism rates were tested using the limma package.

### Codon adaptation index (cAI)

The codon adaptation index was calculated for a given sample as described by Sharp & Li (ref 1987). Briefly, for each sample, a set of the top 10% expressed genes was defined using batch-corrected TPM; the relative synonymous codon usage was then calculated, dividing the observed frequency of each codon by the frequency expected assuming all synonymous codons for a given amino acid are used equally; the codon adaptation index (cAI) is then calculated by comparing the frequency of each codon to the frequency of the most abundant (or optimal) codon for a given amino acid. All codes used to perform this analysis are available on GitHub (https://github.com/elisadonnard/CodonOPT). Codon cAI and gene cAI scores were calculated for both brain cortex and neurons, which are largely identical (data not shown). Only neuron cAI’s and gene cAI scores are presented.

### Codon-stability coefficient (CSC) analysis

CSCs were calculated as previously described (23, 24, 26, 27). Specifically, a Pearson’s correlation coefficient was calculated for each of the 60 non-start and -stop codons between the frequencies of this codon in all the genes that use this codon, and the stability of these genes. The stability of a gene (y), which is the inverse of its degradation rate (x), is expressed as follows:

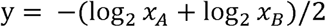

Here *x*_*A*_ and *x*_*B*_ are the normalized degradation rates from library A and B, respectively. The highest expressed isoform of each gene was used to calculate the usage frequencies of each codon.

### Minimum free energy (MFE) analysis

The sequences of 5’UTR, CDS, and 3’UTR were first extracted from the highest expressing isoform of each gene using Biostrings (v2.54.0) (68), minimum free energy (MFE) of each sequence was then calculated using RNALfold module in ViennaRNA (v2.4.14) (69) with the default parameters.

### RNA Immunoprecipitation

HEK 293T cells (ATCC) were cultured with DMEM media, supplied with 10% FBS and 1% Antibiotic Antimycotic Solution (Gibco). Cells of passages between 6 and 9 were transfected with lipofectamine 3000 based on manufacture’s instruction in 6-well plate. 48 hr post transfection, cells are quickly washed in ice cold PBS buffer and collected in complete homogenization buffer as used for ribosome profiling above, plus 1% NP40. After incubating on ice for 10 min, lysate is cleared by centrifuging at 20,000 rcf for 10 min at 4°C. 10% of the lysate was collected as input sample, the rest of the lysate is diluted 1:10 with NT2 buffer (50 mM Tris-HCl (pH 7.4), 150 mM NaCl, 1 mM MgCl_2_, 0.05% NP40) (70). Each half of diluted lysate is incubated with 1/10 volume of antibody coated Protein G Dynabeads™ (Invitrogen) for 40 min at 4°C by end to end rotation. The beads have been coated with Monoclonal ANTI-FLAG® M2 antibody (Sigma-Aldrich) or unimmuned mouse IgG (Santa Cruz). After washing 3 times with NT2 buffer, the beads and the input sample are extracted with TRIzol™ Reagent (Invitrogen), and RNA is precipitated overnight. cDNA is made with QuantiTect Reverse Transcription Kit (Qiagen). qPCR is performed to quantify the reporters (27) with mCherry sequences, and Fmr1 gene, with primers mCherry-F gtacgggtcaaaagcttacgtt, mCherry-R ccttctggaaatgaaagtttaagg, Fmr1-F cgcggtcctggatatacttc, Fmr1-R tggagctaatgaccaatcactg. Recovery rate of the reporters and of Fmr1 is calculated as the ratio of immunoprecipitated RNA versus the input RNA. To account for differences in immunoprecipitation efficiencies between experiments, normalized recovery rate is calculated as the ratio of the recovery rate of the reporter versus the recovery rate of *Fmr1* in each technical replicate, using the recovery rate of *Fmr1* as an internal control.

## Code availability

All codes used to perform cAI analysis are available on GitHub (https://github.com/elisadonnard/CodonOPT). Other customized R scripts for data analysis are available from the corresponding authors upon request.

## Data availability

The data supporting the findings of this study have been deposited in the Gene Expression Omnibus (GEO) repository with the accession code GSE140565 and GSE140642. All other data are available from the corresponding authors upon reasonable request.

## Acknowledgements

We thank Emmiliano Ricci for sharing his experience and protocol for ribosome profiling, Mariya Ivshina for advice and help with the mouse breeding. We thank Lindsay Romo for sharing her protocol for 5-EU labeling of neuron culture, and Jeff Coller (Case Western Reserve University) for Click-iT EU tagging – RNA-seq protocol. We also thank Ariel Bazzini (Stowers Institute for Medical Research) for his very helpful discussion and input. Nathan Gioacchini and Yongjin Lee helped preparing yeast and fly spike-in RNA samples. This works was supported by the National Institutes of Health (U54HD82013); the Simons Foundation, and the Charles H. Hood Foundation (to JDR).

## Author information

### Contributions

H.S. and J.D.R. conceived the project and designed the experiments. H.S. performed most of the experiments, B.L. generated RNA-seq libraries. H.S., E.D. and R.W. performed the bioinformatic analysis. H.S. and J.D.R. wrote the manuscript with input from all authors.

### Corresponding authors

Correspondence to Huan Shu or Joel Richter.

## Figure legends

**Figure S1:**
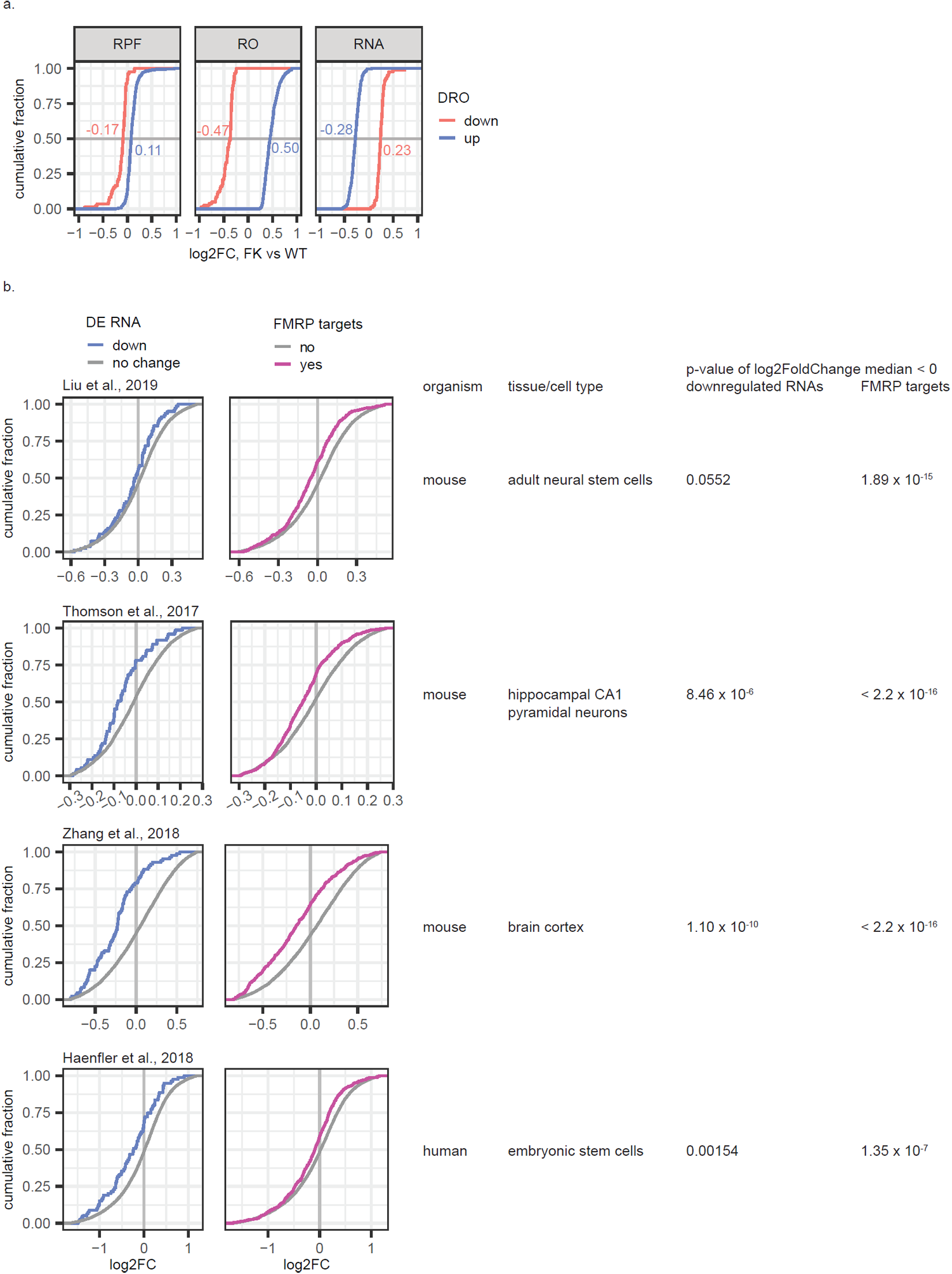
RNA levels changes in Fragile X mouse and human models. **a**, ECDF (empirical cumulative distribution function) plots showing log2FC of RPF, RO (ribosome occupancy), and RNA levels for mRNAs having statistical significant different ROs as calculated using Xtail (31) between FK and WT (2 biological replicates for each genotype, FDR < 0.05). Numbers show the means of each group. **b**, ECDF (empirical cumulative distribution function) plots for log2FoldChange in published RNA-seq data sets of various FXS models (20, 47, 48, 71) for FMRP binding targets (5) (purple; left) and mRNAs downregulated in FK brain cortex in current study (blue, right). The animal species and tissue/cell typed used in each of these studies is indicated. P-values were calculated for the log2FoldChange values of the FMRP targets (purple) and of the downregulated mRNAs identified in this study (blue) to be smaller than 0 (Wilcoxon test, lower tail). For data from Thomson et al. (48), genes were filtered for normalized counts between 10^2.5^ and 10^4.25^ as was done in the original publication. For data generated using human embryonic stem cells (71), only genes with unique mouse orthologs were considered.

**Figure S2:**
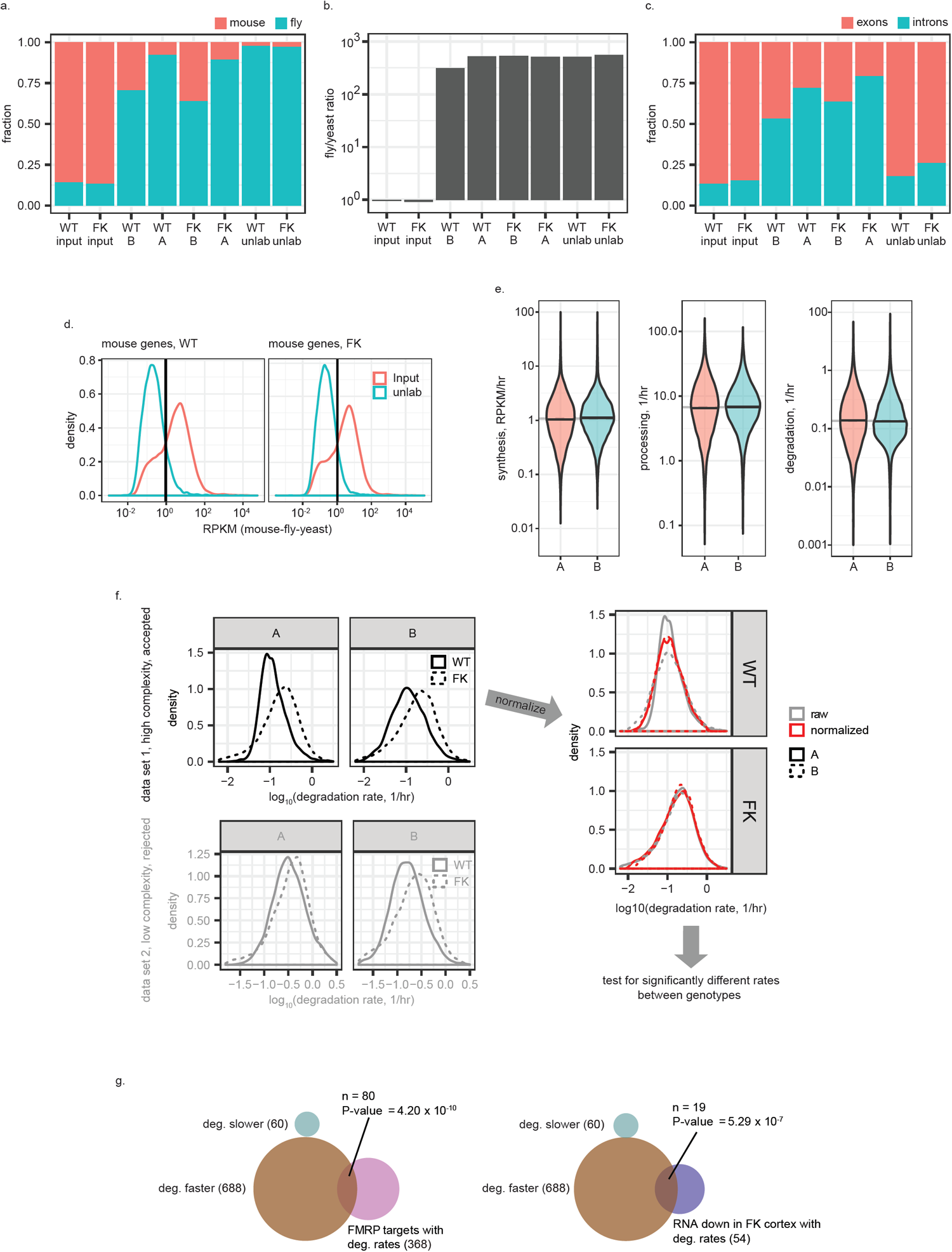
RNA metabolic profiling in WT and FK neurons. **a**, Bar graph of fractions of reads uniquely mapped to mouse (orange) or fly (*Drosophila*) (green) transcriptome in each library. As expected, the pulled-down libraries (A, B and unlab) were enriched in reads mapped to the fly transcriptome (5-EU labeled to saturation) over that of mouse (5-EU labeled only for a brief pulse). The input libraries were not subjected to pull-down and had more reads mapped to mouse compared to fly. Unlab libraries had the smallest ratio of reads that mapped to mouse, demonstrating minimum background to the pull-down process. Accordingly, libraries from mouse neurons that are labeled for a shorter time (20min, libraries A) had smaller ratios of reads mapped to mouse transcriptome than that labeled for longer (60min, libraries B). **b**, Bar graph of ratio of reads that uniquely mapped to *Drosophila* transcriptome *vs* that to yeast in each library. Ratios are scaled so that the mean of this ratio in WT input and in FK input libraries is 1. Similar to panel **a**, the high *Drosophila* to yeast ratio demonstrates specific pull-down to enrich for 5-EU labeled RNA. **c**, Bar graph of fractions of reads that uniquely mapped to exons (orange) and introns (green) among those uniquely mapped to the mouse transcriptome. As expected, input libraries are composed mostly of mature mRNAs and therefore had predominantly exon reads. Similarly, the exon/intron ratio for unlab libraries represents nonspecific signal that originates from the input RNA pool. Libraries from mouse neuron RNAs that are labeled for short (20min, A) or longer (60min, B) are mostly composed of nascent transcripts and therefore had more introns. Accordingly libraries labeled for a shorter time (A) had more introns than that labeled for longer (B). **d**, Density plots of RPKM (read per kb per million reads uniquely mapped to mouse-fly-yeast combined genome) of each mouse gene in input (orange) and unlab (green) libraries in WT (left) and FK (right) neurons. Filtering thresholds (black vertical lines) were identified for WT and FK at 0.95 and 1.05 RPKM, respectively. Genes were filtered for those that had RPKM higher than threshold in input (i.e., that are expressed) and lower than threshold in unlab libraries (i.e., that do not have high nonspecific pull-down background). Data of genes that survive filtering in both WT and FK libraries are analyzed by the INSPEcT program (37) to estimate RNA metabolism rates. **e**, Violin plots for synthesis, processing and degradation rates estimated for libraries A and B. Each violin contains data from both WT and FK. Black horizontal lines denote median of each violin. Grey horizontal lines denote the median of both violins. **f**, Pipeline for normalization and statistical tests for genes with differential RNA metabolic rates, using the degradation rate as the example that is shown. We observed reproducible global faster degradation rates in FK than WT in both libraries A and B and in both data set 1 (high quality libraries, presented in this study) and data set 2 (independent data set, low complexity lacks statistical power and was rejected for gene-level analysis (gray shaded graphs) but was sufficient for confirming global-level shift) (left; **Table S4**). To capture the global shift between genotypes while testing for genes with significantly different metabolism (i.e., synthesis, processing, and degradation) rates, we considered library A and B in data set 1 as pseudo-replicates and normalized them using the Limma package (67) for each genotype separately. With normalized RNA metabolic rates, genes with significantly different rates between genotypes were then called (right). **g**, Venn diagrams showing overlaps between mRNAs with faster (brown) or slower (green) degradation rates and FMRP CLIP targets (left, pink). Numbers of mRNAs in each group and each overlap, as well as the p-value of enrichment of each overlap, are indicated.

**Figure S3:**
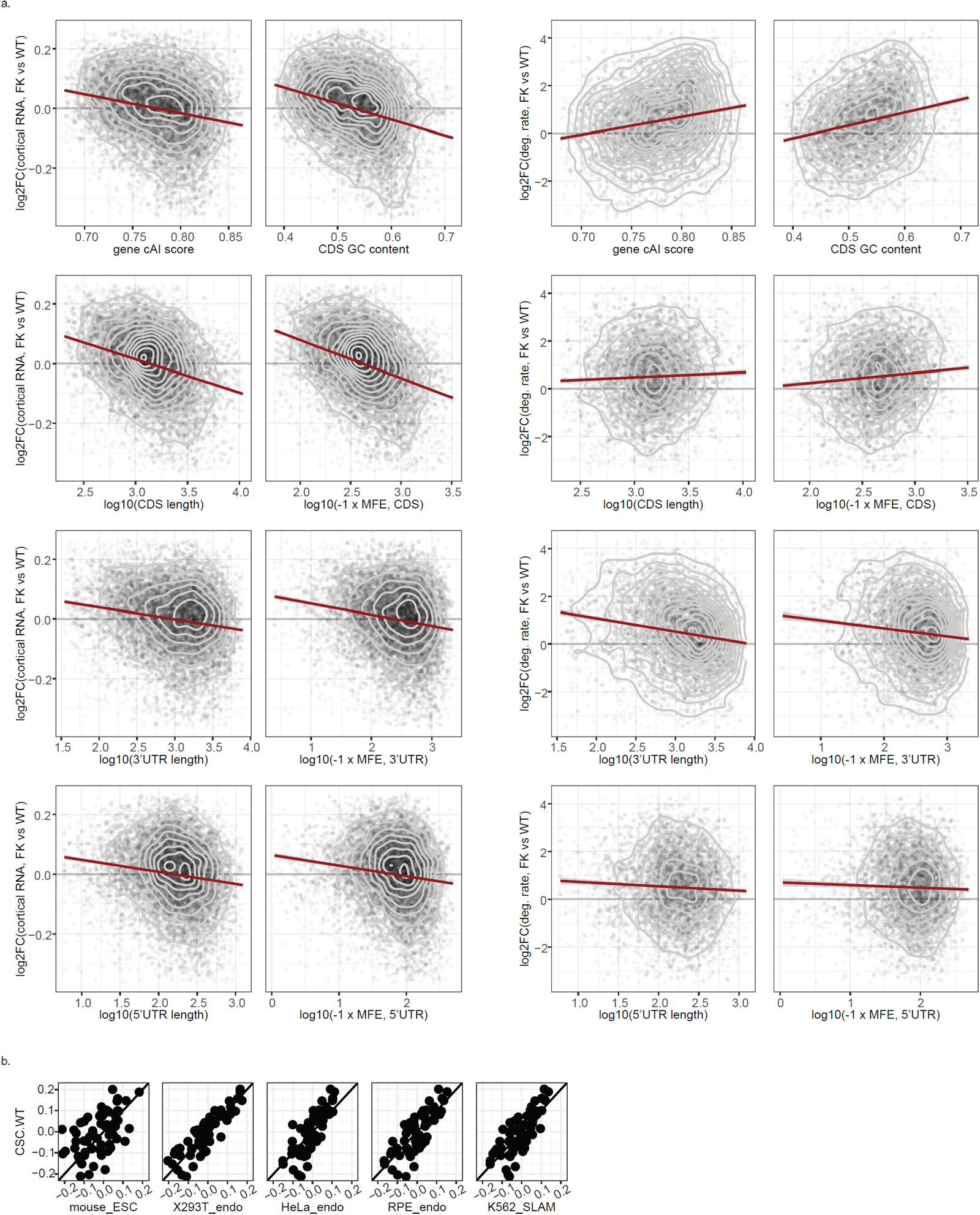
RNA changes and codon optimality. **a**, Scatter and 2D density contour plots of log2FC of steady state RNA levels in FK cortex versus WT (left) and that of degradation rate in FK neurons versus WT (right) as a function of a series of features of the transcripts as indicated. The highest expressed isoform of each gene was used. The red straight line shows the linear regression of the data points. Pearson’s product−moment correlation coefficients are indicated. **b**, Scatter plots comparing CSC (codon stability coefficient) of all non-start or -stop codons in WT neurons in this study (y axis) and that in several mouse and human cell lines shown by Wu et al (27). CSCs in mouse ESC were calculated by Wu et al. based on data published by Herzog et al (72). CSCs in human cells lines (293T, HeLa, and RPE) measured by either blocking transcription (“endo”) or SLAM-seq (“SLAM”) were generated by Wu et al (27).

**Figure S4:**
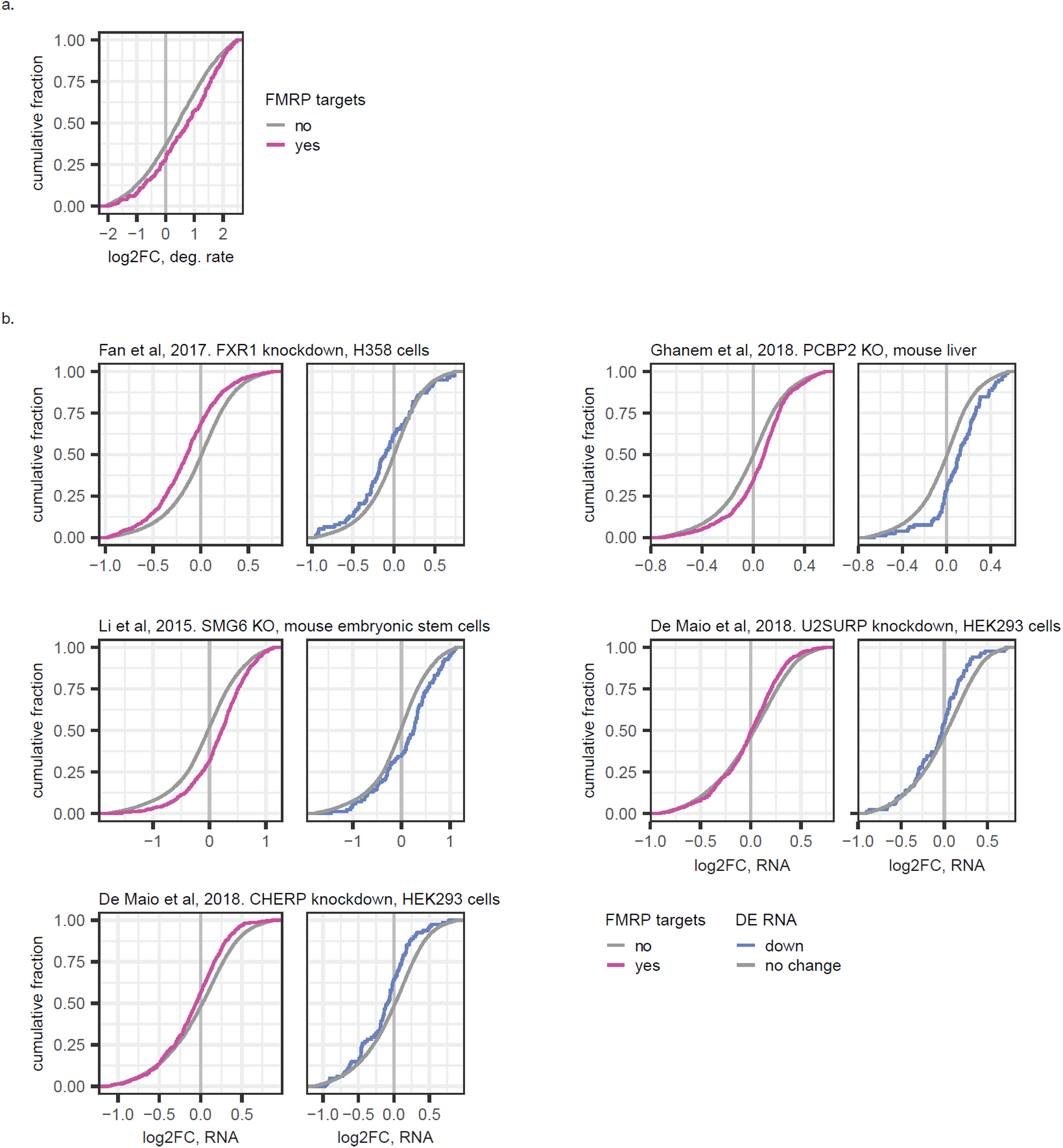
Additional protein factors that may participate in FMRP regulated optimal codon dependent mRNA stability. **a**, EDCF plot showing log2FC of degradation rate change in FK neurons versus WT for FMRP binding targets and non-targets. **b**, ECDF plots showing log2FC of RNA levels for FMRP binding targets (purple) and mRNAs downregulated in FK cortex (blue) after depletion of FXR1 (45), SMG6 (73), PCBP2 (74), U2SURP and CHERP (75). For the dataset generated using human cell lines (45, 75), only genes with unique mouse orthologs were considered.

**Figure S5:**
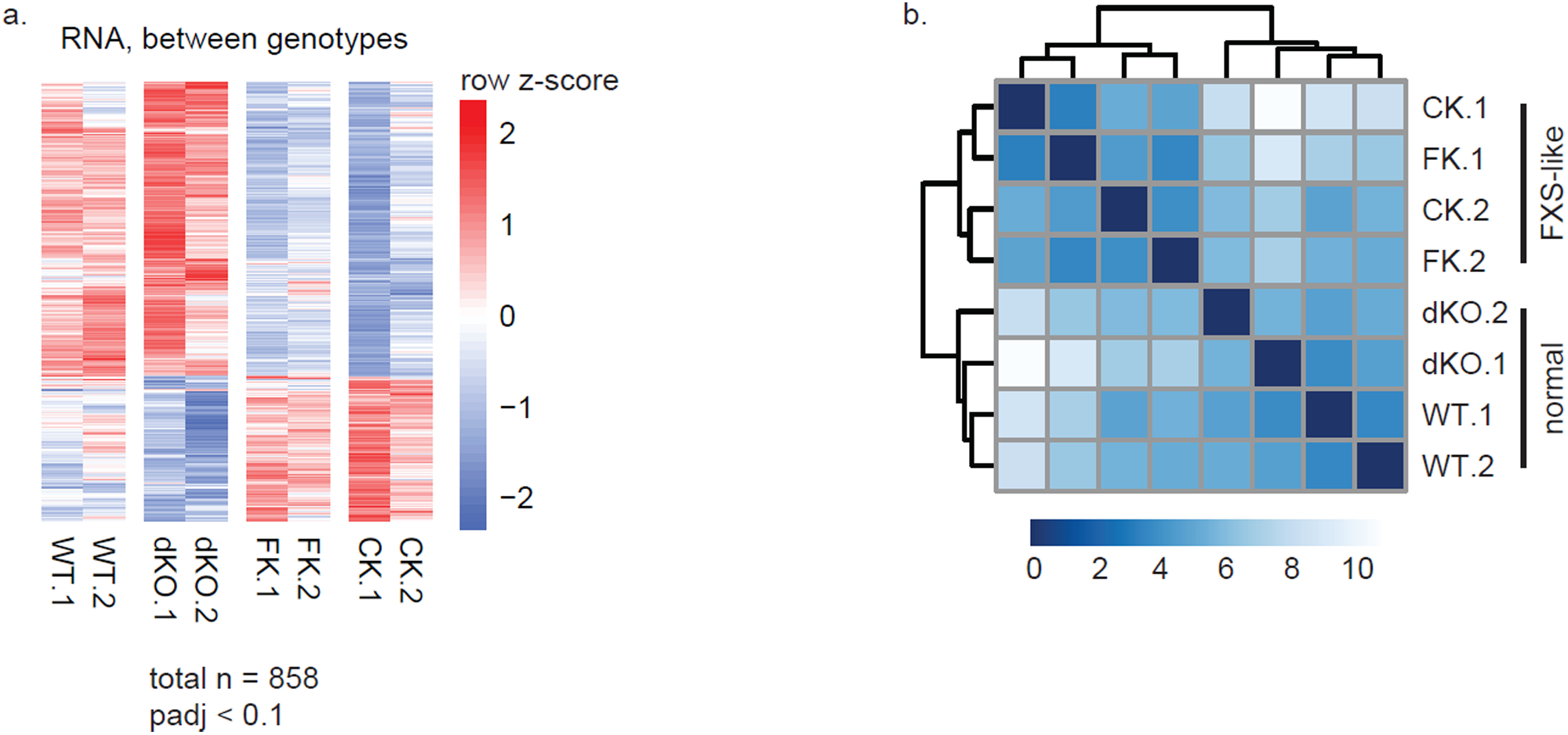
RNA levels changes among the four genotypes of the FXS rescue paradigm. **a**, Heatmap showing differentially expressed mRNAs between any two genotypes of the four genotypes (WT, FK, CK, and dKO). Red and blues shades show high or low z-scores for each mRNA (row) across all samples. Both replicates are plotted separately for each genotype. **b**, Unsupervised hierarchical clustering of sample to sample distances measured by the Euclidean distance between each other using their top 1000 most variable mRNAs. Darker to lighter shades of blue indicate closer to farther distance between samples. Dendrogram represents the clustering.

**Figure S6:**
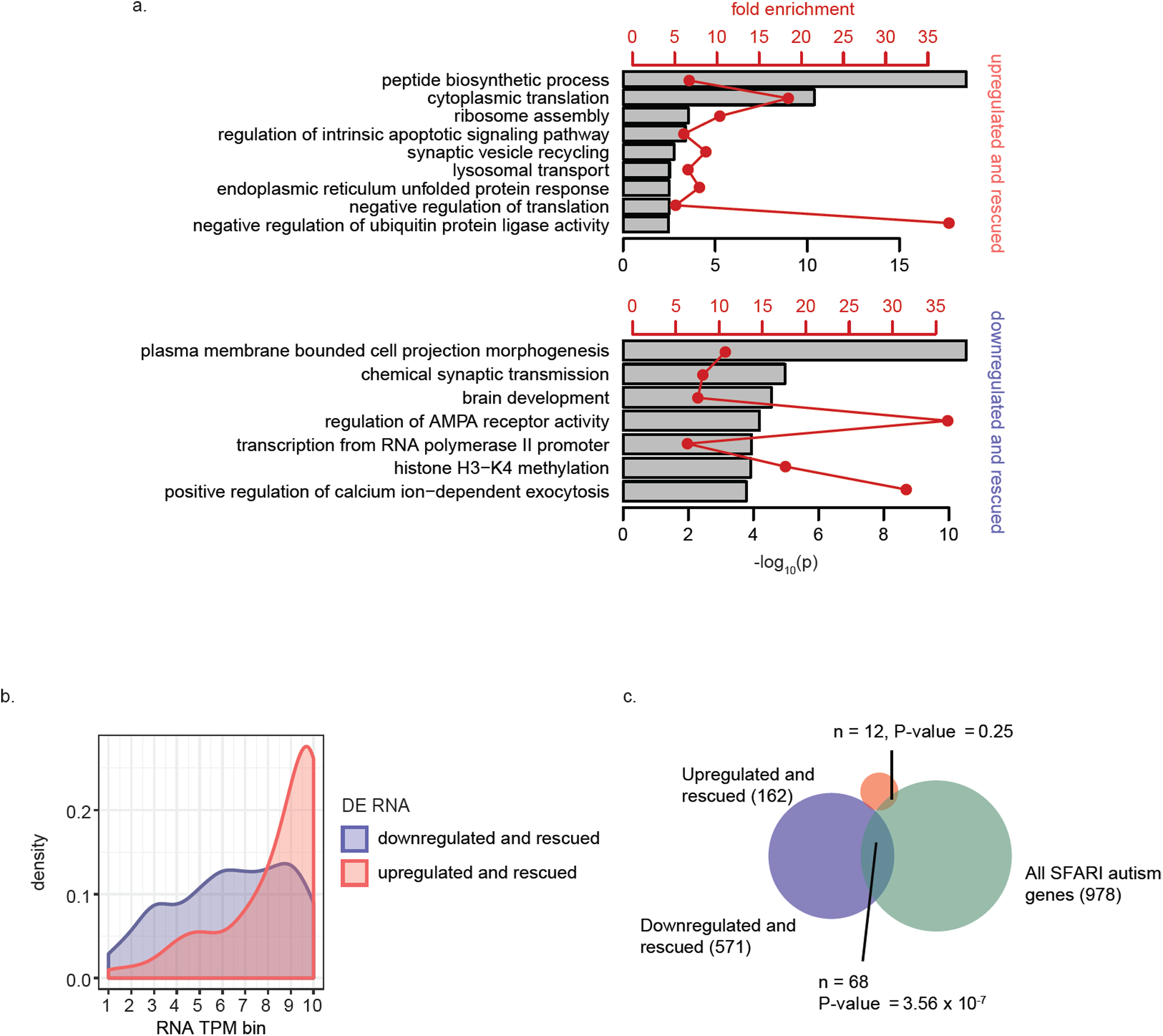
Additional information about the mRNAs differentially expressed between the normal (WT and dKO) and the FX-like (FK and CK) groups. **a**, Representative Gene Ontology (GO) terms enriched for genes upregulated (upper) or down regulated (lower) at the RNA level in the FXS-like group. Grey bars and red point-and-lines show the –log_10_(P value) and fold enrichment of each of these GO terms, respectively. See **Table S2-3** for full lists of enriched GO terms. **b**, Density plot of the distribution of mRNAs up- (red) or down- (blue) regulated in the FXS-like group over RNA transcript per million (TPM) bins. Bins were generated by dividing all detectable protein coding genes into 10 equal bins based on their TPM in WT brain. Bin 1 genes have TMPs of the lowest quantile and bin 10 the highest quantile. **c**, Venn diagram showing the overlap between the differentially expressed mRNA with all SFARI autism risk genes (51). Numbers of mRNAs in each group and in each overlap as well as p-values of enrichment (hypergeometric test, upper tail) are indicated.

